# High-throughput screening of aqueous two-phase systems for polyphenol extraction from *A. nodosum*: a green chemistry approach

**DOI:** 10.1101/2021.06.28.450208

**Authors:** Alex Olivares-Molina, Brenda Parker

**Affiliations:** The Advanced Centre for Biochemical Engineering, Department of Biochemical Engineering, University College London, London, UK

**Keywords:** Aqueous two-phase system, Automation, Biorefinery, Green chemistry, Polyphenol recovery

## Abstract

Brown macroalgae are an attractive third-generation feedstock of natural products, in order to design green chemistry-compliant processes and reduce the use of organic solvents in bioactive product extraction, aqueous two-phase systems (ATPS) was applied. This research aimed to develop a high-throughput screening (HTS) to recover polyphenols from *Ascophyllum nodosum* using ATPS. In total, 384 different 2-phase systems were assessed using an automated liquid-handling system to evaluate polyphenol recovery using a model system of phloroglucinol to establish an optimal 2-phase system for polyphenol partitioning. Various ratios of PEG:potassium phosphate solutions were explored to evaluate partitioning of polyphenols via a scale-down approach. Scale-down selected system showed a recovery of phloroglucinol of 62.9±12.0%, this system was used for scale-up trials. Scale-up studies confirmed that the HTS method was able to recover polyphenols with a 54.8±14.2% in the phloroglucinol model system. When the optimised ATPS system was tested with a polyphenol extract, 93.62±8.24% recovery was observed. When ATPS was applied to a fucoidan and alginate biorefinery residue, 88.40±4.59% polyphenol was recovered. These findings confirm that ATPS is a valuable addition to the bioprocess toolkit for sustainable extraction of natural products from macroalgae in a multiproduct biorefinery approach.

**Practical application:** Selection of the best concentrations of phase-forming components and recovery conditions for the application of aqueous two-phase systems in an industrial setup has been proved to be laborious and cumbersome. This paper presents an automated platform to rapidly assess several ATPS to recover polyphenols from brown macroalgae and a subsequent confirmation with the scale-up of the potential candidates and contrasted with two case studies. This methodology allows a quick screening for the best aqueous two-phase system and can be expanded to recover high-value products from other types of macroalgae or microalgae.

## 1 Introduction

Current developments in the third generation biorefinery industry are transitioning towards more environmentally sustainable processes, given the need to reduce the environmental footprint of their operations, and the consideration of additional dimensions besides the technical feasibility of a new technique. The transition of promising technologies from bench-scale to commercial level is often difficult and expensive, such as aqueous two-phase systems (ATPS); though in ATPS the high concentrations of salts and the cost of the polymers used are reasons claimed to be the limitation for the industrial application [1], thus scale-down approaches have appeared in recent years to analyse these systems in a more resource-efficient way. ATPS are governed by their phase diagrams and can be considered as a unique fingerprint of a two-phase system under a given set of conditions, namely pH, temperature, or salt concentration. Information then can be derived from such diagrams which includes: the concentration of the phase-forming components necessary to form an ATPS in equilibrium, the subsequent concentration of phase components in the top and bottom phases, and the ratio of phase volumes [2]. Embedded in the phase diagram is the binodal curve, which divides a region of component concentrations that will form immiscible aqueous two-phases, located above the curve, from those that will form one homogeneous phase, at and below the curve. The determination of the binodal curve can be calculated and fitted by least squares regression to the empirical relationship developed by Merchuk et al. (Eq. 1) [3], where *y* and *x* are the concentrations of polymer and salt respectively, which had been determined via experimentation, and *A*, *B*, and *C* which are the regression parameters.

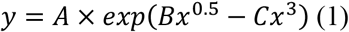

There are no reports of the application of ATPS for the recovery of high-value products from brown macroalgae, although there are publications in the recovery of R-phycoerythrin from red seaweeds [4–6], and one report recovering sulphated polysaccharides from the green seaweed *Enteromorpha* sp. [7] hence, these compounds together with other high-value products are interesting options to evaluate the use of ATPS to extract them from brown algae. Polyphenols from seaweeds can be extracted by different methods, i.e. solvents [8], microwave-assisted extraction [9], enzyme-assisted extraction [10], supercritical fluid extraction [11], among others; the main drawback of these methods is either economic scale-up and/or environmental performance. Recovery of polyphenols from macroalgae using ATPS have not been reported until the submission of this document, but there are reports on the recovery of polyphenolic compounds from lignocellulosic material [12, 13], thus is interesting to explore the extraction of these compounds applying green chemistry strategies, summarised in the use of ATPS for their recovery.

The mechanism of separation in ATPS is complex and not easy to predict [14]. In general, the partition of high-value products is a result of Van der Waals and ionic interactions of the natural compounds with the surrounding phase. Some of the key mechanisms have been identified: electrostatic, hydrophobic, and steric hindrance interactions between the phases and the compound of interest. The most important factors that influence ATPS are those inherent to the phases, and in PEG/salt based ATPS, increases in salt concentration result in an increase in partition coefficients of bioproducts in the upper phase or interface because of salting out. Another important factor involved in the performance of ATPS despite the composition of the phases is pH. Partitioning of compounds into the phases in an ATPS is determined by their isoelectric point (pI). The pH of the system, however, affects the charge of the wanted compound and ion composition as well as introduces differential partitioning into the two phases. Consequently, the initial pH of the system must be above the pI of target bioactives [15]. Considering a conventional polymer-salt aqueous two-phase systems for industrial production, the substances and equipment needed for its preparation are rather inexpensive compared to initial capital expenditures of previously mentioned methods, and recycling of phase-forming components is simpler than recycling or disposal of solvents [16]. Also, is important to consider that limitations in the processing speed, equipment intricacy and thus scalability concerns are minimal since simple apparatus like stirred tanks or disc stack centrifuges, and mixer-settlers can be used [1]. Unluckily, the apparent technical simplicity of the setup of the process is not paralleled by the complex and still unresolved parameter predictability for an industrial purification step. When there is a gap in the knowledge about a certain molecule or process, the experimental approach in terms of high-throughput process development is a solution for a broad exploration of the design space. High-throughput process development is a methodology for the use of large number of parallelised, miniaturised, and automated experiments [17]. High-throughput technologies can be used for characterisation of aqueous two-phase systems for the determination of binodal curves for different salts and polymers [16]. In ATPS, the first step to consider, as previously explained, is the determination of the binodal curve, but the process to determine or estimate this curve is cumbersome and require extensive analytical work and time, thus a rapid evaluation is needed. Reports in the purification of protein compounds, antibodies, pDNA among others have been described, yet a high-throughput procedure for polyphenols has not been defined yet [18–20]. Since theoretical or model based solutions for this dilemma have not been developed, the only practical solution lies in a rapid and automated screening approach allowing parameter evaluation of a multi-parameter space within a short period of time. The method of choice for such a screening approach is found in the use of high throughput screening (HTS) platforms. These platforms are already used on a routine basis in the field of high throughput analytics. HTS applications in the development of ATPS and downstream process development in general have received attention in recent years [17, 21, 22], but it has never been used for the extraction of polyphenols from brown seaweeds.

In this study, the development of a methodology for a rapid evaluation of aqueous two-phase systems for polyphenol recovery using an automated high-throughput screening approach was used for the preparation, characterisation and application of 384 different polymer-salt ATPS using a commercially available pipetting robot TECAN Freedom Evo™ 200. Preparation was simply established by liquid pipetting in combination of adequate mixing of the two-phase systems. Characterisation of the obtained systems was carried out by an automated determination of binodal curves and further estimation of regression parameters. Finally, polyphenol partitioning had been investigated, first by rapid proxy quantification of a model molecule for polyphenols, and later with scale-up trials of selected two-phase systems using two complex polyphenolic samples from brown seaweeds. The approach presented in this chapter is based on a screening only varying potassium phosphate buffer and polymer concentrations, being able to process close to 770 samples per day, and it offers a rapid screening method for the study of the partitioning behaviour of high-value compounds from macroalgae with potential economic breakthrough using ATPS.

### 2 Materials and Methods

### 2.1 Reagents and materials

Analytical grade Folin-Ciocalteu reagent, sodium carbonate (Na_2_CO_3_), sodium phosphate monobasic and dibasic, phloroglucinol, polyethylene glycol (PEG) 1000, 2000, 3000 and 6000 were purchased from Sigma-Aldrich (Saint Louis, Missouri, USA). Analytical grade methyl orange dye was from VWR International (Radnor, Pennsylvania, USA). A Millipore milli-Q (Burlington, Massachusetts, USA) water purification system was used to purify the water used in all experiments. *A. nodosum* was purchased from Hebridean Seaweeds Company Ltd (Isle of Lewis, UK), where it was wild harvested and dried and milled to a particle size of 300 μm. All macroalgal material was kept dry and stored in plastic containers at room temperature to proceed with the subsequent experiments.

### 2.2 Apparatus

High-throughput screening (HTS) experiments were performed using a Freedom Evo™ 200 (Tecan, Crailsheim, Germany) automated liquid handling station (LHS). This LHS was used to prepare all aqueous two-phase systems (ATPS), binodal determination, and phloroglucinol partitioning. Freedom Evo™ 200 was equipped with an 8-channel air liquid handling (LiHa) arm with conductive polypropylene disposable tips connected to 0.5 mL dilutors. This station was controlled by Evoware Standard 2.6™ software.

### 2.3 Optimisation strategy for the extraction of polyphenols

The strategy used to optimise ATPS partitioning followed a scale-down approach combined with HTS. The determination of the binodal curves was performed using 10 μL methyl orange dye, a hydrophobic compound that will always partition to the top phase. 2% w/w screening variation of the phase-forming components was used to build the said curves.

The manual preparation of phase systems was conducted in 1.5 mL micro-centrifuge tubes. Appropriate amounts of PEG solutions were added, followed by potassium phosphate buffer, water, and methyl orange dye. The systems were mixed using a vortex for 15 s, followed by a settling time of 1.5 h. All systems were built to a working volume of 800 μL. Pipettes were used for sampling the bottom phase. All experiments were conducted at 23±2°C.

Completed the manual determination, and prior to the automated determination of the curves pipetting was calibrated by adjusting the relevant parameters such as aspiration and dispensing speed and height, air gap sizes and waiting times for each solution to be pipetted and a 2% w/w screening variation of the phase-forming components was used. Phase systems were prepared according to Bensch et al. [16] with minor modifications; subsequence addition of PEG 1000, 2000, 3000, and 6000, phosphate buffer and water in 1.2 mL 96-deep-well plates. For the determination of binodal curves a 10 μL of methyl orange was added into each sample well. All systems reached a total volume of 800 μL and were mixed through repeated aspiration and dispensing cycles and allowed to separate for 1.5 h. After the settling time was completed, 50 μL were aspirated from the bottom of the deep-well plate and transferred into a microplate. Finally, separation of phases was observed and phase-forming systems were it occurred were selected together with the one-phase systems preceding them (without phase separation) to build the binodal curves estimating the parameters of the exponential fit (Eq. 1) developed by Merchuk et al. [3] using the curve fitting toolbox on MATLAB.

After the estimation of the binodal curves screening, the partition of polyphenols was assessed using HTS with the LHS and phloroglucinol as a model molecule. All the systems assessed were built on the same manner as explained in the automated determination of the curves, but a fixed amount of phloroglucinol was added to each ATPS (10 μL; 0.25 mg/mL) instead of methyl orange. All the systems were allowed to partition for 1.5 hour, and after completed this step 50 μL were aspirated from the top phase of the deep-well plate and transferred to a microplate for absorbance quantification at 267 nm.

The best candidate ATPS mixtures to partition polyphenols were identified and further scale-up of these systems was evaluated. Systems were scaled up 40 times completing a total volume of 50 mL and the recovery yield between the scaled-down and scaled-up systems was quantified according to the following section.

### 2.4 Soluble polyphenol extraction

Polyphenol extraction was completed according to Olivares-Molina & Fernández (2016) with minor modifications. Firstly, 1 g of dried and milled macroalgae was weighed and placed into glass shaking flasks. An aliquot of 50 mL of acetone:milli-Q water 70:30 (v/v) was added, and samples were incubated for 1 h with continuous shaking at 250 rpm in an ISF-1-V Climo-Shaker Incubator and Platform Shaker (Kuhner, Switzerland) at room temperature. This step was repeated four times, and between extractions samples were centrifuged at 3200×*g* for 6 minutes in an Avanti J-E centrifuge (Beckman Coulter, UK). Supernatants of each individual sample were collected in a single container, to produce one individual sample of each replicate. After extraction acetone was removed by evaporation in a Genevac EZ-2 centrifugal evaporator (SP Scientific, USA). The aqueous extracts were centrifuged for 15 minutes at 3,200×*g* to remove residual biomass, and the supernatant then was vacuum-filtered through 0.7 μm pore GF/F filters (Whatman, UK), and frozen for subsequent analyses.

### 2.5 Total polyphenol quantification

The total polyphenol content cannot be quantified in presence of non-ionic detergents, i.e. PEG, due to the precipitation of the Folin-Ciocalteu reagent [24]. To be able to quantify the total polyphenolic content a surfactant was added and the original method developed by Jerez et al. [25] was modified accordingly. 100 μL of polyphenol extract was mixed with 500 μL of Folin-Ciocalteu reagent (diluted 1:10 v/v) and 20 μL of sodium dodecyl sulphate (SDS), this mix was homogenised using a vortex, and allowed to stand 3 minutes at room temperature. After this, samples were centrifuged at 14000×*g* for 10 min and supernatant was transferred to a new micro-centrifuge tube. Next, 400 μL of 7.5% w/v Na_2_CO_3_ was added to the mixture, homogenised, and incubated for 15 minutes at 45°C in a Thermomixer R 2.0 mL block (Eppendorf, Germany). Following the incubation 200 μL of every sample were placed in a 96-well plate and the absorbance was measured at 765 nm in an Infinite M200 PRO microplate reader (Tecan, Switzerland). Blank samples were prepared using milli-Q water instead of polyphenol extract. Total polyphenol content was expressed as micrograms of phloroglucinol equivalent per mg of dried alga (μg PGE/mg alga).

### 2.6 Statistical analysis

Data were expressed as the means ± standard deviations (SD). For group comparisons the Student t-test for independent samples was performed where applicable (p<0.05). When analysis of variance (ANOVA) was needed the distribution of grouped variables was calculated first to determine the homoscedasticity of data. In addition, to check if data was parametric, Cochran’s C and Levene’s tests were applied (p>0.05). After data homogeneity was confirmed, a one-way ANOVA was performed to verify if significant differences (p<0.05) occurred between the samples. Parametric data was analysed using the post-hoc Tukey HSD test (p<0.05). All statistical analyses were performed in STATISTICA 7.0 suite.

## 3 Results and Discussion

ATPS can generally be characterised by the determination of the binodal curve, and the respective partition coefficient for a given system. On the condition that all these parameters are known for various ATPS a rapid evaluation of their feasibility in a potential bio-purification process becomes possible. The strategies followed to reach this goal are summarised in Fig 1.

**Fig 1.**
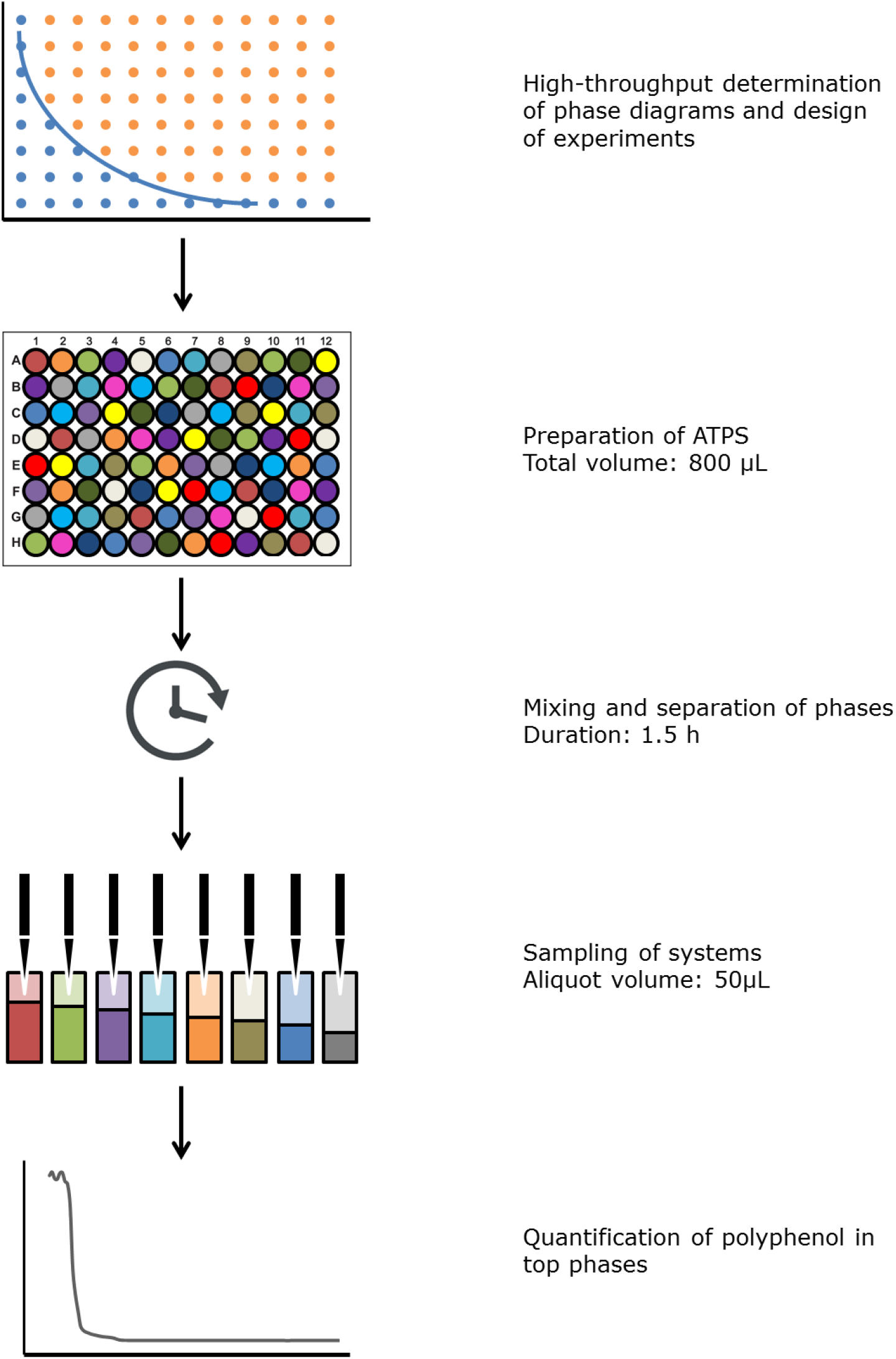
Overview of the developed high-throughput screening platform. After phase diagrams were determined in high-throughput, ATPS were selected for the partitioning screening. ATPS were prepared in deep-well plates, and phloroglucinol was added from a stock solution. After mixing and settling, samples were taken from the top phase, and polyphenols were quantified through absorbance.

**Fig 2.**
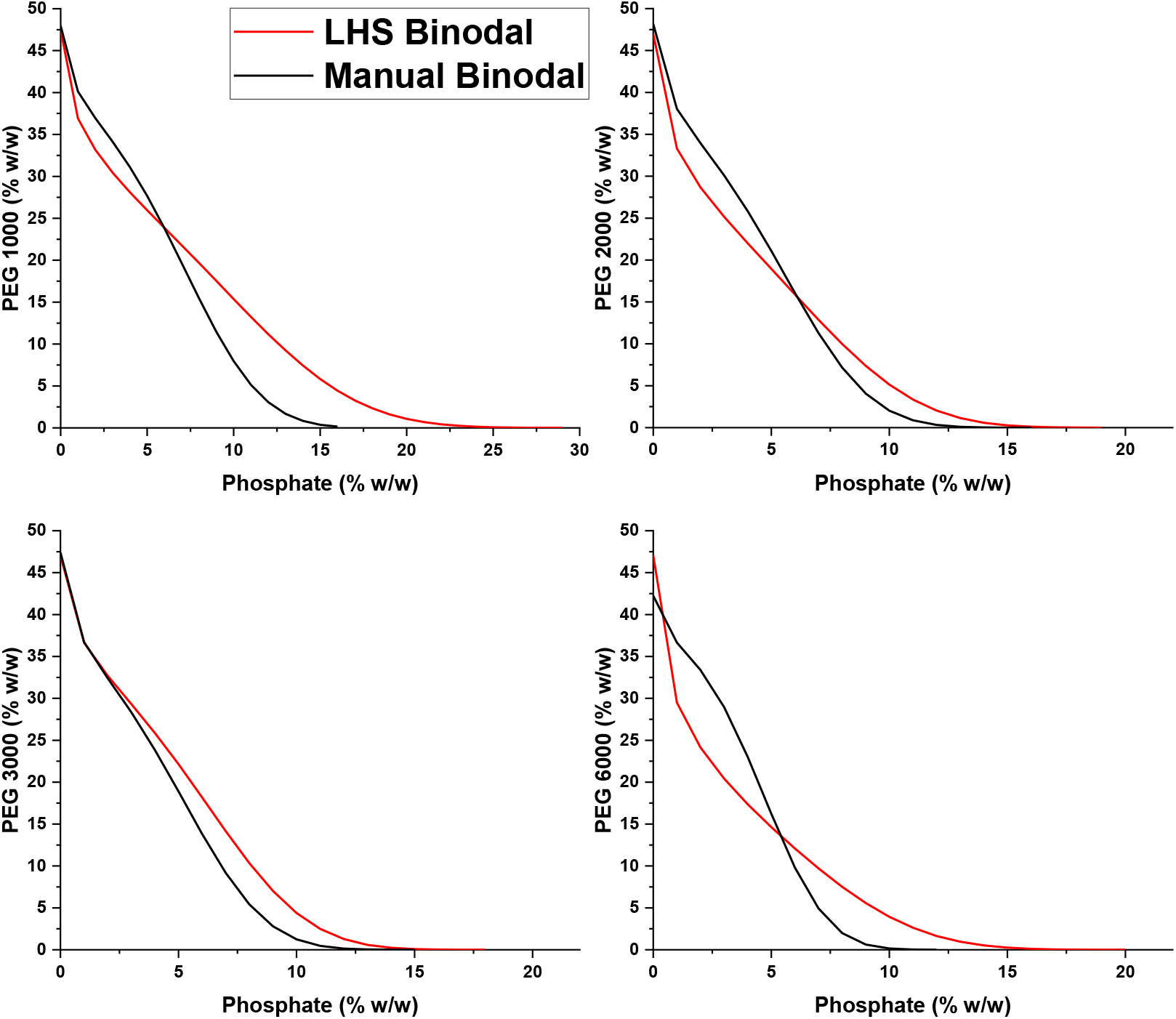
Comparison of binodal curves determination of 4 phase-forming systems: (A) PEG 1000, (B) 2000, (C) 3000, and (D) 6000; with phosphate buffer using an automated method and a manual determination process.

### 3.1 Method development and validation

Prior to its use, the method was optimised in terms of sample volume, ease of handling, accuracy, and reproducibility. Generally, the most convenient way of mixing on liquid handling stations lies in the use of orbital shakers. However, many applications of orbital shaking systems reported are hampered by insufficient power input, low shaking orbit, or non-optimised parameters [20]. For this reason, mixing process on the robotic system was conducted in deep well plates where disposable tips were used for aspirating and re-dispensing 800 μL of the phase system. Samples were taken after seven aspiration/dispensing cycles [16]. Aqueous two-phase systems are firstly characterised by determining the binodal curve, which is the boundary condition for the development of the two-phase systems between the polymer and the salts involved. For the systems investigated in this study, methyl orange always completely partition to the PEG-rich phase. In one-phase systems, methyl orange is distributed homogenously over the entire system. In two-phase systems on the other hand, the dye partitioned completely to the top phase leading to an almost transparent bottom phase.

### 3.2 Determination of binodal curves

To validate the high throughput methodology, ATPS were determined both manually and by use of the liquid handling system. The determination of the binodal curve represents the boundary conditions for the formation of a two-phase system, and is a necessary prerequisite for any ATPS partitioning study. When it is done manually, these determinations are performed using a turbidimetric titration assay or by analysing the PEG/salt concentrations in the top and bottom phase of a series of two-phase systems. Both methods are difficult to adapt to an automated environment or require high analytical effort, e.g. labour, reagents, and time. To determine the formation of the two-phase systems a hydrophilic dye was used, methyl orange, that always partition to the top phase and in one-phase systems the dye is distributed homogenously over the entire system [27]; therefore, the sample aspirated from the bottom phase showed an orange colouration. Binodals were determined for the PEG-potassium phosphate systems by use of the LHS. The experimental parameter space was initially covered by a matrix of 96 PEG-potassium phosphate mixtures, as the position of the binodal is unknown a rather wide matrix was used, a sampling space of 4% w/w of phase-forming components between points was selected. A regression fitting using Merchuk’s equation (Eq. 1) has shown accurate goodness of fit in previous reports [20, 28]. The estimation parameters for the binodal line and their 95% confidence bounds with the LHS are displayed in Table 1 and the R^2^ of the fit for the curves were, PEG 1000-potassium phosphate: 0.9802, PEG 2000-potassium phosphate: 0.9904; PEG 3000-potassium phosphate: 0.9902, ad PEG 6000-potassium phosphate: 0.9777.

**Table 1.** Regression coefficients of binodal curves estimation built using the liquid handling station method and their 95% confidence bounds.

| | Phase components | A ($\pm 95\%$ CI) | B ($\pm 95\%$ CI) | C ($\pm 95\%$ CI) |
| --- | --- | --- | --- | --- |
| Automated method | PEG 1000-Phosphate | 47.48 (44.12, 50.83) | -0.25 (-0.30, -0.20) | $-3.3 \times 10^{-4}$ ($-4.6 \times 10^{-4}$, $-2.1 \times 10^{-4}$) |
| | PEG 2000-Phosphate | 47 (44.06, 49.94) | -0.34 (-0.41, -0.27) | $-1.1 \times 10^{-3}$ ($-1.7 \times 10^{-3}$, $-6.1 \times 10^{-4}$) |
| | PEG 3000-Phosphate | 47.03 (43.98, 50.07) | -0.25 (-0.32, -0.18) | $-1.6 \times 10^{-3}$ ($-2.2 \times 10^{-3}$, $-9.9 \times 10^{-4}$) |
| | PEG 6000-Phosphate | 47.08 (43.18, 50.98) | -0.47 (-0.58, -0.36) | $-1.0 \times 10^{-3}$ ($-1.8 \times 10^{-3}$, $-1.8 \times 10^{-4}$) |
| Manual method | PEG 1000-Phosphate | 47.95 (40.39, 55.51) | -0.18 (-0.29, -0.065) | $-1.2 \times 10^{-3}$ ($-1.9 \times 10^{-3}$, $-5.2 \times 10^{-4}$) |
| | PEG 2000-Phosphate | 48.17 (42.25, 54.1) | -0.23 (-0.35, -0.12) | $-2.4 \times 10^{-3}$ ($-4.1 \times 10^{-3}$, $-7.1 \times 10^{-4}$) |
| | PEG 3000-Phosphate | 47.41 (44.34, 50.49) | -0.25 (-0.32, -0.19) | $-2.8 \times 10^{-3}$ ($-3.6 \times 10^{-3}$, $-2.0 \times 10^{-3}$) |
| | PEG 6000-Phosphate | 42.24 (34.93, 49.55) | -0.14 (-0.30, -0.027) | $-5.2 \times 10^{-3}$ ($-8.7 \times 10^{-3}$, $-1.7 \times 10^{-3}$) |

**Table 2.** Error between manual and automated determinations of the binodal curve.

| Systems | Error (%) |
| --- | --- |
| PEG 1000-Phosphate | 2.09±0.29 |
| PEG 2000-Phosphate | 1.47±0.30 |
| PEG 3000-Phosphate | 1.26±0.28 |
| PEG 6000-Phosphate | 2.14±0.27 |

After the automated determination of the binodal curves was completed a manual determination was performed, in order to compare both procedures and validate the robustness of the LHS approach. The experimental parameter space, as well as the LHS approach, was covered by a matrix of 96 PEG-potassium phosphate mixtures, and similarly than the LHS method a sampling space of 4% w/w of phase-forming components between points was selected to assess the transition of a one- and two-phase system. The estimated parameters of Eq. 1 from the manual determination with their 95% confidence bounds are exhibited in Table 1. Goodness of fit of the binodal curves from the manual determination exhibited an R^2^ around 0.80 – 0.99 for the manual method (Fig 3), with PEG 1000-potassium phosphate showing a R^2^ = 0.8856, PEG 2000-potassium phosphate R^2^ = 0.953, PEG 3000-potassium phosphate R^2^ = 0.9847 and PEG 6000-potassium phosphate R^2^ = 0.7998, being more variable than the automated determination.

**Fig 3.**
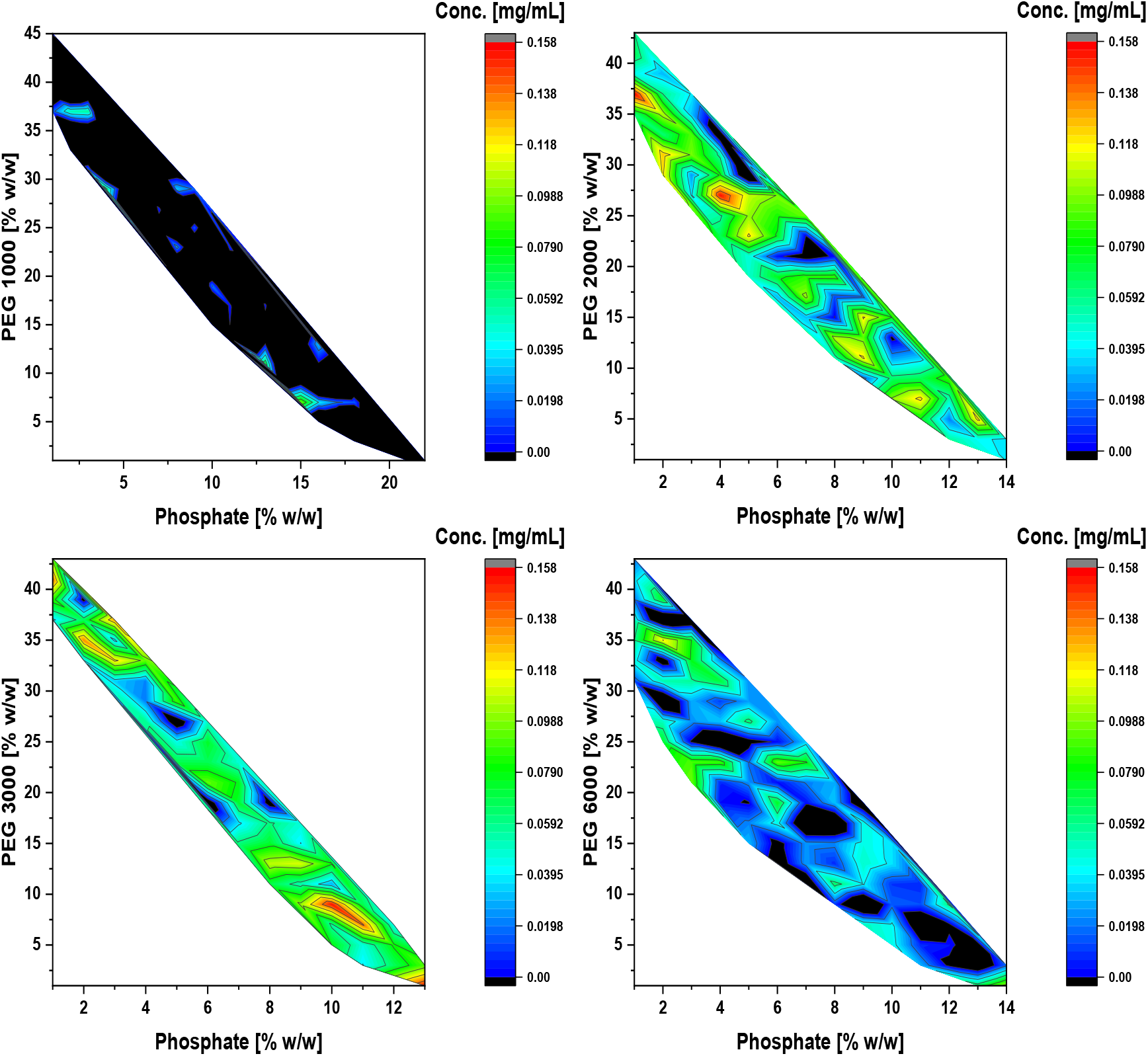
Contour plots of the partitioning of phloroglucinol in ATPS formed with A) PEG 1000, B) 2000, C) 3000, and D) 6000; with phosphate buffer.

Completed both, manual and automated determinations, the methods were compared between them. All regression fittings showed binodal data similar with phase systems published previously, except from Huddleston et al. [29]: Huddleston et al. have worked with PEG 1000-K_3_PO_4_ systems at 25°C, Bensch et al. [16] have worked with PEG 200, 1000, 1540, and 4000 with potassium phosphate buffer systems at 25°C, and Lei et al. [30] have worked with PEG 400, 600, 1000, 1500, 3400, 8000, and 20000 and potassium phosphate systems at 4°C instead of 25°C and at pH 7 instead of 7.5. The values obtained by Huddleston et al. differ from the values obtained in the current work; this may be due to the fact that they did not use a buffered solution but only added K_3_PO_4_, a salt which is expected to yield a much higher pH value than the 7.5 used within this work. The difference between manual and automated determination was between ~1.25 – 2.15% with the lowest value in the PEG 3000-potassium phosphate systems and the highest with PEG 6000-potassium phosphate systems (Table 3). The binodal fitting of all the automated approach curves exhibited a goodness of fit within the 95% confidence limits of the averaged fitted curve. The most accurate automated fitting was in the PEG 3000-potassium phosphate system, with an error in the comparison of 1.26±0.28%; it can be noted as well that the automated binodal in each point of the regression is above the manual binodal, overlapping this curve. This situation was desirable because any error between the methods is negligible due to any two-phase system formed above the actual binodal curve is contained in the LHS determination. Following, PEG 2000-potassium phosphate systems displayed an error between the regressions of 1.47%±0.30%, while PEG 1000-potassium phosphate, and PEG 6000-potassium phosphate systems binodal regressions showed an error of 2.09±0.29% and 2.14±0.27%, respectively. Differently from PEG 3000-potassium phosphate system, in all the other curve comparisons there was an intersection point between the regression approaches located in the Cartesian plane in (6, 15.99) for PEG 2000-potassium phosphate, (6, 23.84) for PEG 1000-potassium phosphate, and (6, 10.95) in PEG 6000-potassium phosphate system, this means there was an over- and underlapping in different portions of the fitting of the automated method with the manual approach. These slight deviations might arise from incorrect readings when performing the titration manually or from a too-high distance between two system points evaluated using the hydrophobic dye marker, and thus leading to a deviation when performing the regression. The accuracy of the latter can be enhanced by choosing a smaller step value by a second run focusing on the binodal region [16]. Nevertheless, from a processing perspective, both the experimental parameter space and accuracy shown in Fig 3 can be considered as sufficient since system points close to the binodal curve do not express the necessary process robustness needed in industrial purification procedures. This outcome confirmed that automated construction of the binodal curve is equally reliable than manual determination and is a suitable method to assess polyphenol partitioning.

### 3.3 High-throughput screening of polyphenol partitioning

After the validation of the automated screening method was demonstrated, a high-throughput screening for partitioning of polyphenols was performed. The above methodology represents a tool for the characterisation of system properties describing the respective ATPS and thus prepares the stage for the actual polyphenol partitioning. The LHS was used to make 384 different ATPS and subsequently analyse the recovery of polyphenols. For the preparation of the phase systems generally four pipetting steps for water, buffer, a fixed concentration of phloroglucinol of 0.25 [mg/mL] and PEG solution were required (see Fig 1). Phloroglucinol was selected as a model polyphenol to assess the partitioning due to participation as the main building block in the synthesis of polyphenols in brown macroalgae [26]. Analysis using direct absorbance at 265 nm required one additional pipetting step for sampling of the top phase, and transfer into a 96-well microplate. In order to reach a statistical robustness in the determination of the polyphenol partitioning, nine independent samplings were performed for each system, the measured absorption values when using the direct absorbance method showed an overall standard error of the mean (SEM) of: SEM = 6.8 [μg/mL] for PEG 1000-potassium phosphate systems, SEM = 5.2 [μg/mL] for PEG 2000-potassium phosphate, SEM = 5.6 [μg/mL] for PEG 3000-potassium phosphate, and SEM = 3.8 [μg/mL] for PEG 6000-potassium phosphate. This suggests both, a high precision during the pipetting steps, as well as the accuracy of the overall assay. The first results that can be observed in Fig 3 is that mostly all the systems constructed with PEG 1000-potassium phosphate buffer did not partition phloroglucinol. This might be due to the shorter hydrophilic end group with shorter polymer chains that reduces the hydrophobicity of the top phase [31]. Similarly, PEG 6000-potassium phosphate buffer systems did not recover phloroglucinol as well; a possible explanation of this effect might be due to the steric hindrance of PEG 6000 with water, not allowing the transport of phloroglucinol to the top phase, as well as the fact that PEG with higher molecular weights produce smaller coefficient factor between top and bottom phases [32]. On the other hand, in PEG 2000-potassium phosphate buffer systems it was observed areas where the polyphenol partitioning occurred, highlighted in red colouration in the contour plots, in two areas with the highest partitioning in PEG2000-potassium phosphate (37:1) and (27:4), with 0.157±0.030 [mg/mL] and 0.155±0.066 [mg/mL], respectively; and the total recovery yield of both systems was 62.9±12.0% and 61.8±26.6%, respectively. Finally, in PEG 3000-potassium phosphate buffer systems areas with high partitioning were also observed in the contour plot, in PEG 3000-potassium phosphate (9:10) and (1:13), with 0.154±0.070 [mg/mL] and 0.153±0.050 [mg/mL], respectively; and 61.6±27.9% and 61.2±19.7%, respectively. From these four prospective recovery systems just PEG 2000-potassium phosphate buffer (27:4) and PEG 3000-potassium phosphate buffer (9:10) were selected for further scale-up trials; this decision was due to PEG 2000-potassium phosphate buffer (37:1) and PEG 3000-potassium phosphate buffer (1:13) system points were close to the binodal curve regression limit, thus the associated error with the formation of the two phases was higher and the extraction process cannot be considered robust enough to use it in an industrial setup.

### 3.4 Scale-up trials of selected ATPS

The selected ATPS from the partitioning screening were scaled-up 40 times and compared with the scaled-down systems to validate the process. As stated above PEG 2000-potassium phosphate (27:4) scale-down system exhibited a recovery of 62.9±12.0%, while the partitioning of the scale-up system was 54.8±14.2%. On the other hand, PEG 3000-potassium phosphate (9:10) recovery of the scale-down ATPS was 61.6±27.9% compared to 40.4±14.6% of the scale-up system. While scale-up and scale-down approaches in both systems statistically are not different (Fig 4), there is not a *p* > 0.95 of significance to determine the best scaled-up system. PEG 2000-potassium phosphate (27:4) exhibited a significance of *p* > 0.70 compared to a significance value of *p* > 0.25 of PEG 3000-potassium phosphate (9:10); thus, PEG 2000-potassium phosphate (27:4) exhibited a better statistical significance and therefore was selected for the final tests with two complex samples.

**Fig 4.**
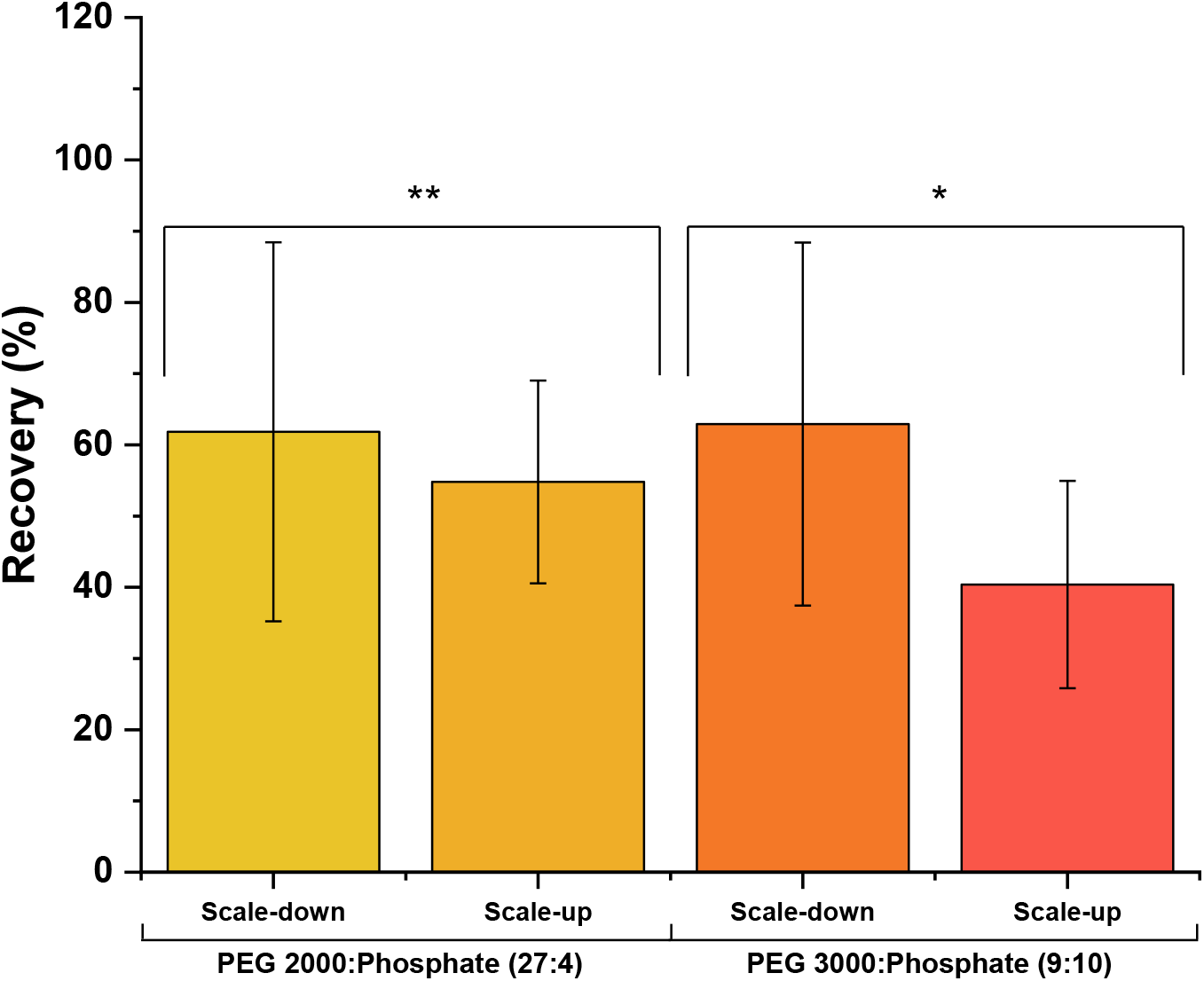
Comparison of the recovery between selected scale-down systems (800 μL) and scale-up tests (50 mL). Values grouped with “**” showed a significance of p > 0.70, and values grouped under “*” showed a significance of p > 0.25.

Thus, PEG 2000-potassium phosphate (27:4) was selected to test two case studies to confirm the optimisation of the processing step, a polyphenol partitioning trial from an aqueous polyphenol extract from *A. nodosum*, and another test recovering polyphenols from a biorefinery residue that extracted fucoidan and alginate first. The recovery of the aqueous extract was 93.6±8.24%, and the recovery of the biorefinery residue was 88.4±4.59%. When compared with the few results that exists in literature PEG 2000-potassium phosphate (27:4) exhibited a 10-fold increase in the partitioning, with 11.38±1.00 [g PGE/100 g DW alga] and 10.76±0.56 [g PGE/100 g DW alga] for polyphenol extract and biorefinery residue respectively, while Xavier et al. [12] showed a partitioning yield of 1.88±0.04 [mg GAE/100 g DW wood] using a PEG 2000-ammonium sulphate (1.08% w/w) system from eucalyptus wood industrial wastes. Both partitioning trials did not showed statistical significant differences (*p* < 0.05) between them, however both experiments exhibited differences with the phloroglucinol model (Fig 5), demonstrating a better performance with complex samples and polyphenol samples with more than one polyphenol type, this effect might be due to the isoelectric point (pI) of the polyphenols within the raw extract and biorefinery residue are closer to pI of the ATPS (pI = ~7.5 – 8.5). These results confirm that PEG 2000-potassium phosphate (27:4) is the optimal ATPS for polyphenol partitioning from *A. nodosum*.

**Fig 5.**
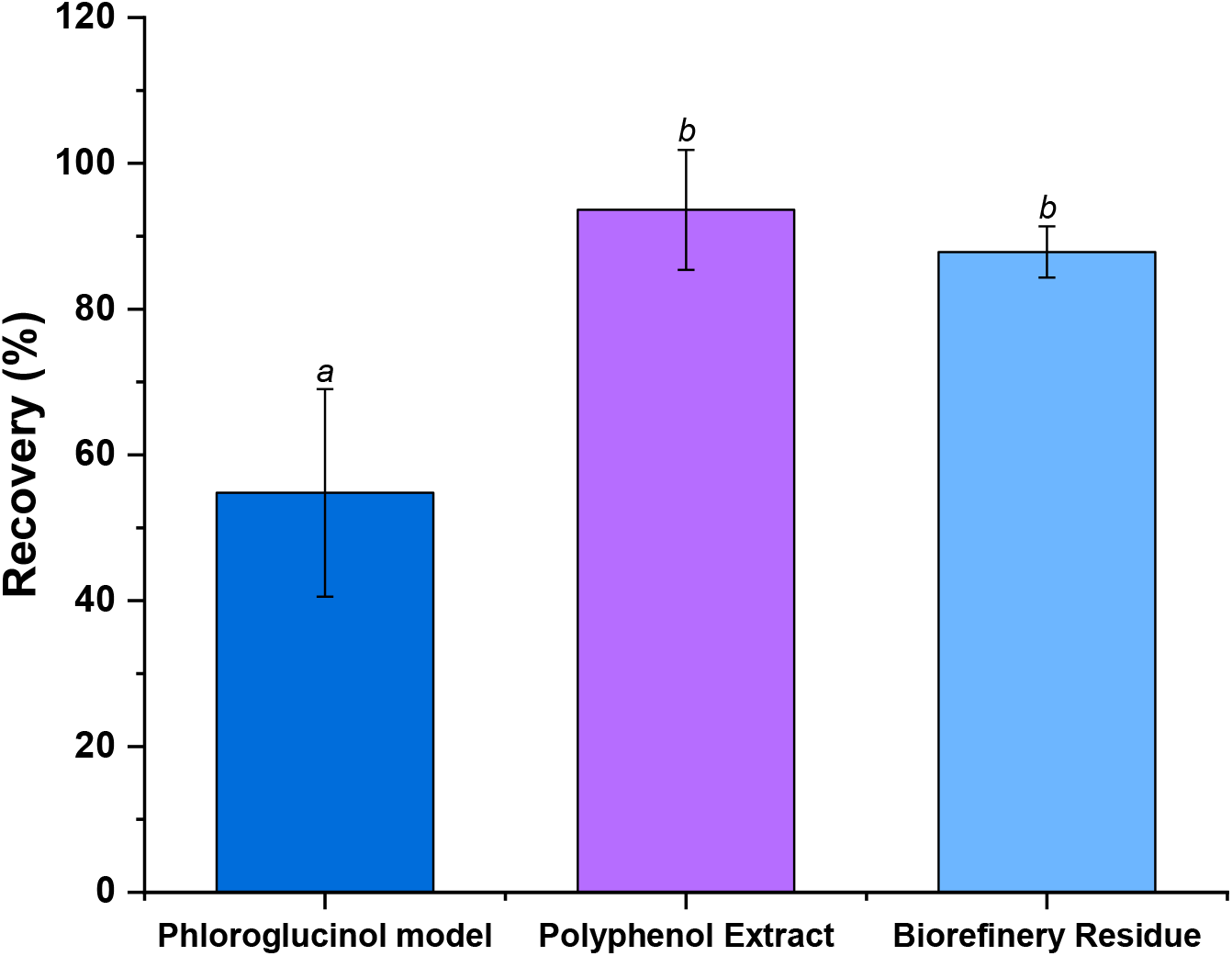
Comparison of the recovery of the scale-up model and two case studies: a polyphenol extract, and a carbohydrate biorefinery residue. Values are means ± standard deviation (n=3). Values with different letters are significantly different, as determined by a post hoc Tukey’s HSD test (*p* < 0.05).

The benefit of the use of the high-throughput approach presented in this publication is that resource intensive experiments and/or potential new processing steps, such as application of aqueous two-phase system for polar compounds extraction, can be instead assessed in a scale-down manner to see whether a separation is feasible based on process performance requirements. In addition, this methodology allows the exploration of process robustness issues, based on a fast and reliable assessment of the binodal curve determination. The main limitation of this approach is that the method is a high-throughput assessment that allows the possibility to design ATPS processes while screening a broad range of parameters, by only varying the concentration of the phase-forming components but does not take into account any process optimisation regarding to temperature or different settling times.

## 4 Concluding remarks

The main aim of this work was to develop a platform for high-throughput screening experimentation to select a suitable aqueous two-phase system extraction to separate polyphenols from its macroalgal matrix. The use of a high-throughput screening approach was chosen so that a rapid and robust methodology for process development could be obtained. Partitioning of polyphenols was investigated with four different polymers, PEG 1000, 2000, 3000, and 6000 using an automated liquid handling station in a scale down approach. Binodal curves for every system were built with the automated system and compared with a manual determination, proving being equally reliable than a manual binodal curve determination. Partition of polyphenols was affected by polymer molecular weight, where the smallest and largest weights (PEG 1000 and 6000) did not partition polyphenols, while intermediate weights (PEG 2000 and 3000) exhibited 62.9±12.0% and 61.6±27.9% recovery values in the scale-down method. Finally, during the scale up trials PEG 2000-potassium phosphate buffer (27:4) showed the highest recovery of all 384 systems analysed with 93.62±8.24% and 88.40±4.59% of polyphenols in the top polymer-rich phase recovered from a polyphenol extract and a biorefinery residue.

## Abbreviations

ATPS: aqueous two-phase system
HTS: high-throughput screening; PEG, polyethylene glycol
PGE: phloroglucinol equivalent
pI: isoelectric point
LHS: liquid-handling station
SEM: standard error of the mean

Alex Olivares-Molina was supported by a “Becas Chile” scholarship granted by the National Agency of Research (ANID), Chile.

The authors have declared no conflict of interest.

